# Efficacy of a *Pseudomonas aeruginosa* Serogroup O9 Vaccine

**DOI:** 10.1101/2023.07.13.548830

**Authors:** Dina A. Moustafa, Antonio DiGiandomenico, Vishnu Raghuram, Marc Schulman, Jennifer M. Scarff, Michael R. Davis, John J. Varga, Charles R. Dean, Joanna B. Goldberg

## Abstract

There are currently no approved vaccines against the opportunistic pathogen *Pseudomonas aeruginosa*. Among vaccine targets, the lipopolysaccharide (LPS) O antigen of *P. aeruginosa* is the most immunodominant protective candidate. There are twenty different O antigens composed of different repeat sugars structures conferring serogroup specificity, and ten are found most frequently in infection. Thus, one approach to combat infection by *P. aeruginosa* could be to generate immunity with a vaccine cocktail that includes all these serogroups. Serogroup O9 is one of the ten serogroups commonly found in infection, but it has never been developed into a vaccine, likely due, in part, to the acid labile nature of the O9 polysaccharide. Our laboratory has previously shown that intranasal administration of an attenuated *Salmonella* strain expressing the *P. aeruginosa* serogroup O11 LPS O antigen was effective in clearing and preventing mortality in mice following intranasal challenge with serogroup O11 *P. aeruginosa*. Consequently, we set out to develop a *P*. *aeruginosa* serogroup O9 vaccine using a similar approach. Here we show that *Salmonella* expressing serogroup O9 triggered an antibody-mediated immune response following intranasal administration to mice and that it conferred protection from *P. aeruginosa* serogroup O9 in a murine model of acute pneumonia.

## INTRODUCTION

*Pseudomonas aeruginosa* is a Gram-negative bacterium and an important opportunistic pathogen. It is one of the primary bacteria responsible for nosocomial infections; in particular, acute infections leading to sepsis in patients with ventilator-associated pneumonia and those with burn wounds, surgical incisions, diabetic foot ulcers, and catheters. It is also the primary cause of chronic lung infections in individuals living with cystic fibrosis, leading to morbidity and mortality in this population. In other patients, acute lung infections can lead to chronic infections. *P. aeruginosa* can also infect otherwise healthy individuals, causing otitis externa, otitis media, folliculitis, and keratitis; most of these infections are due to a breach of normal host immune defense (1, 2).

Infections with *P. aeruginosa* are of particular concern because this bacterium is naturally antibiotic resistant and becoming multidrug-resistant (MDR) or extensively drug resistant (XDR). MDR *P. aeruginosa* is considered a “Serious Threat”, as defined by the CDC Antibiotic Resistance Threats in the United States Report-2019. And *P. aeruginosa* infections are notoriously difficult to treat once established. Thus, there is an urgent need to develop new methods to combat infections caused by *P. aeruginosa*.

Vaccination represents an appropriate approach to prevent infections by *P. aeruginosa*, however, vaccines targeting this bacterium have been under investigation for over 50 years, with none having yet been approved (3–6). The surface-exposed lipopolysaccharide (LPS) is the immunodominant protective antigen of *P. aeruginosa* and therefore is considered an appropriate target for vaccine development. LPS is composed of the lipid A embedded in the outer membrane, the core oligosaccharide, and O antigen polysaccharide, which extends out from the surface of the bacterial cell. For *P. aeruginosa*, twenty different International Antigenic Typing System (IATS) serogroups are recognized, based on the expression of the O antigen portion. All of the O antigen structures have been determined (7) with each serogroup possessing subtype strains having subtle variations, leading to over 30 subtypes (8). The serogroup-specificity of the O antigen suggests that a comprehensive *P. aeruginosa* LPS-based vaccine would need to encompass all these subtypes. Fortunately, numerous studies have found 10 serogroups to be most common in various types of infection (9–12).

One attractive approach would be to develop a cocktail of the most common LPS serogroups as a vaccine. However polysaccharides are generally considered poor immunogens and alone do not elicit a robust immune response (reviewed in (13)). Because of this, many of the currently available polysaccharide based-vaccines are polysaccharide-protein conjugates (14). For *P. aeruginosa*, in the 1980s-1990s the Swiss Serum and Vaccine Institute developed and tested an octavalent conjugated vaccine with eight different *P. aeruginosa* serogroups covalently coupled to the exotoxin A antigen of *P. aeruginosa* (15). While the clinical results of these studies were never reported, this vaccine did not contain serogroups O8 or O9, because they contain internal ketosidic linkages (16) and thus cannot be adequately separated from the toxic lipid A component using acid hydrolysis needed to conjugate to the protein carrier. More recently, Nasrin et al. (12) used a “Multiple Antigen Presenting System” based on high molecular weight polysaccharides (17) and have targeted 8 of the most common *P. aeruginosa* O antigen serogroups (12). Serogroup O8 and O9 were also missing from this system.

Consequently, we set out to develop a vaccine to one of these “neglected” O antigen serogroups of *P*. *aeruginosa*. Our laboratory has previously shown that intranasal administration of an attenuated *Salmonella* strain expressing the *Pseudomonas aeruginosa* serogroup O11 LPS O antigen was effective in clearing and preventing mortality in mice following intranasal challenge with serogroup O11 *P. aeruginosa* (18). Here we show that *Salmonella* expressed serogroup O9 and can trigger an antibody-mediated immune response following intranasal administration that conferred protection in a murine model of acute pneumonia.

## MATERIALS AND METHODS

### Cloning and expression of *P. aeruginosa* serogroup O9 O antigen on *S. typhimurium*

Genomic DNA was isolated from *P. aeruginosa* serogroup O9 strain PAO9 (kindly provided by Gerald B. Pier, Harvard Medical School, Boston, MA), using standard procedures. Genomic DNA was randomly sheared through a syringe needle and was end-repaired and cloned into pWEB::TNC (Epicentre Technologies, Madison, WI), followed by packaging into MaxPlax lambda packaging extracts. The lambda particles were used to infect *Escherichia coli* EP105. Colonies were absorbed with mouse monoclonal antibodies to *P. aeruginosa* serogroup O9 (Rougier Bio-Tech Ltd.).

Colonies reacting with antisera were separated with anti-mouse antibodies bound to magnetic beads (Dynabeads; Thermo Fisher Scientific) followed by magnetic bead separation using a mini-magnetic particle separator (CPG Inc. Lincoln, Park, NJ). Positive colonies were selected by colony immunoblot using anti-serogroup O9 mouse monoclonal antibody. The serogroup O9 locus was then cloned from the pWEB::TNC plasmid into the broad host range cosmid vector, pLAFR376 (kindly provided by Laurence Rahme, Massachusetts General Hospital, Boston MA). To do this, plasmid DNA from a positive colony was digested into approximately 20-25 kb fragments with *EcoRl* and ligated to completely digested pLAFR376. The ligation reactions were packaged as bacteriophage lambda particles with the Stratagene Gigapack® XL-11 packaging system and used to infect *E. coli* HB101. Transformed cells were selected on Luria broth (LB) media containing (tetracycline (Tet) 10 μg/ml) where serogroup O9 positive clones were identified by colony immunoblots using the anti-serogroup O9 mouse monoclonal antibody. The positive clone that exhibited the strongest reaction was grown in broth culture for further characterization. This plasmid was isolated and designated pLAFRO9. The construct was confirmed by enzyme digestion and DNA sequencing (Emory Integrated Genomics Core). Plasmid sequences were assembled using PlasmidSPAdes (3.13.1) and the presence of the serogroup O9 locus was confirmed by BLAST against a reference locus accession AF498420.1. Plasmid pLAFRO9 was transferred to *Salmonella enterica* serovar Typhimurium strain SL3261, by P22 transduction using *S. typhimurium* LB5010 as an intermediate host, as described previously (19) and expression was confirmed by silver-stained gel and Western immunoblot of extracted LPS.

### Preparation of bacterial strains used for immunization and infection

*S. typhimurium* SL3261 containing pLAFRO9 (vaccine) or SL3261 containing the cosmid pLARF376 (vector), were used for immunization. All strains were grown overnight in LB supplemented with 10 μg/ml Tet. Both strains were subcultured and grown to an OD_650_ of 0.5. Bacteria were then washed twice and resuspended in phosphate-buffered saline, pH 7.4 (PBS). For infection, *P. aeruginosa* serogroup O9 was grown on Difco^TM^ *Pseudomonas* Isolation Agar (PIA) overnight at 37°C and suspended in PBS to an OD_600_ of 0.5, corresponding to ∼10^9^ colony-forming units (CFU)/mL. Inocula were adjusted spectrophotometrically to obtain the desired dose in a volume of 20 μL. All strains were adjusted spectrophotometrically to obtain the desired immunization or challenge dose.

### LPS extraction, SDS-PAGE, and Western immunoblotting

LPS extracts of *P. aeruginosa* and *Salmonella* organisms were prepared and separated by sodium dodecyl sulfate (SDS)-polyacrylamide gel electrophoresis (PAGE) as described (18) with a Novex X-Cell Surelock minicell system (Invitrogen, Carlsbad, CA). Tris-bis-polyacrylamide gels (12.5%) were cast in 1.0-mm Invitrogen cassettes. After PAGE separation was completed, lysates were electroblotted onto Trans-Blot 0.2-μm-pore-size pure nitrocellulose membranes (Bio-Rad Laboratories, Hercules, CA) by use of a Bio-Rad Mini Trans-Blot electrophoretic transfer cell. Membranes were blocked and then probed with *Pseudomonas* serogroup O9-specific rabbit polyclonal antibodies (Denka Seiken, Tokyo, Japan), followed by incubation with anti-rabbit secondary antibodies conjugated to alkaline phosphatase (Sigma). Reactions were visualized by the addition of Sigma fast 5-bromo-4-chloro-3-indolylphosphate-nitroblue tetrazolium.

### Immunization with *S. typhimurium* SL3261 and *P. aeruginosa* challenge

Use of animals in this study was reviewed and approved by the University of Virginia Institutional Animal Care and Use Committee (IACUC) under protocol number 2844-02-11. All mice were kept under specific pathogen-free conditions, and all guidelines for humane endpoints were strictly followed. All animal experiments were conducted in accordance with the “Public Health Service Policy on Humane Care and Use of Laboratory Animals” by NIH, “Animal Welfare Act and Amendments” by USDA, “Guide for the Care and Use of Laboratory Animals” by National Research Council (NRC). Female BALB/c mice, 5 to 6 weeks old (Harlan Laboratory [Indianapolis, IN] or Jackson Laboratory [Bar Harbor, ME]), were immunized intranasally as described (18). Prior to vaccination, mice were anesthetized by intraperitoneal injection of 0.2 ml of xylazine (1.3 mg/ml) and ketamine (6.7 mg/ml) in 0.9% saline. For intranasal vaccination, mice were given 1 x 10^7^ - 1 x 10^9^ CFU of either SL3261 (pLAFRO9) (vaccine) or SL3261 (pLAFR376) (vector) intranasally in a 20 μl volume (10 μl per nostril). Boosters were performed using the same protocol as the initial vaccination, approximately 14 days after the initial vaccination.

For bacterial challenge, anesthetized mice were given 20 μl of *P. aeruginosa* strain PAO9 prepared as described above. For *in vivo* protection experiments, infected mice were observed over 5 days, and animals that were moribund following infection or in any way appeared to be under acute distress were humanely euthanized and were included as non-survivors in the experimental results.

### Collection of sera

Sera were collected at four weeks post-immunization as described (18). Briefly, blood was obtained by nicking the lateral tail vein of mice; blood sat at room temperature for about 4 hours and was then placed at 4°C overnight. Serum was removed from the red blood cell pellet and was spun at 1,700 rpm for 10 minutes. Samples were stored at −80°C.

### ELISA analysis of *P. aeruginosa* PAO9 lipopolysaccharide (LPS) expression from ***S. typhimurium***

Enzyme-linked immunosorbent assay (ELISA) analysis was performed on sera as described (18). Briefly, 96-well microtiter plates were coated with *P. aeruginosa* strain PAO9, incubated overnight at 4°C, washed with PBS plus 0.05% Tween 20 (PBS-T), blocked with PBS supplemented with 2% bovine serum albumin (PBS-B), and then washed with PBS-T again. Serum samples were serially diluted in PBS-B and 100 μl were placed into each well in PAO9 -coated plates, in duplicate*. Pseudomonas* serogroup O9-specific rabbit polyclonal antibodies were used as positive controls. After overnight incubation at 4°C, the plates were washed three times with PBS-T and air-dried. Secondary antibodies (anti-mouse total IgG, IgG1, IgG2a, IgG2b, IgG3, or IgM conjugated to alkaline phosphatase (Southern Biotechnology Associates, Inc., Birmingham, AL)) diluted 1:5,000 in PBS-B were then added to individual plates and incubated at 37°C for 1 hour. Plates were developed in the dark for 1 hour with 1 mg/ml 4-nitrophenyl phosphate in substrate buffer (24.5 mg MgCl_2_, 48 ml diethanolamine per 500 ml; pH 9.8); development was stopped by adding 50 μl 3 M NaOH. Plates were read using a plate reader at 405 nm. Data were collected using the SOFTmax PRO software (Molecular Devices Corp., Sunnyvale, CA) and then transferred to GraphPad Prism version 6.0 software (GraphPad Software, San Diego, CA) for analysis. For the Ig titer determination, total serum IgG or IgM absorbance readings were adjusted by subtraction of values obtained from the blank, the x-intercept defined the endpoint titer and represented as the reciprocal dilution. IgG subtype quantification for serum samples was based on standard curves that were designed for each antibody isotype by use of GraphPad Prism version 6 software.

### Detection of bacterial loads

For bacterial load quantification, mice were euthanized at 24 hours post-infection and nasal wash and whole lungs were collected aseptically. For the nasal wash, an 18-G catheter was placed at the oropharyngeal opening of the mouse and 1.0 ml of PBS-B was flushed through the nasal passage and collected. Whole lungs were collected from each mouse, weighed, and homogenized in 1 mL of PBS. Tissue homogenates were serially diluted and plated on PIA and CFU determination was made 16 to 18 hours later. Final results were expressed as CFU/ml for nasal washes and CFU/g for lung tissues (18).

### Passive immunization and infection

All animal procedures were conducted according to the guidelines of the Emory University Institutional Animal Care and Use Committee (IACUC), under approved protocol number DAR-2003421-042216BN. The study was carried out in strict accordance with established guidelines and policies at Emory University School of Medicine, and recommendations in the Guide for Care and Use of Laboratory Animals of the National Institute of Health, as well as local, state, and federal laws. For passive immunization, 6 to 8-week-old female BALB/c mice were anesthetized and infected via the intranasal route with ∼1 x 10^9^ CFU/mouse as described. Antisera collected from PBS-, vector-, or vaccine-immunized mice were delivered to the mice (5 μl /nostril) immediately after infection. Mice were euthanized at 24 h post-infection and nasal washes and whole organs were collected aseptically. Nasal wash collection was performed as described. Lung, liver, and spleen tissues were collected from each mouse, weighed, and homogenized in 1 mL of PBS. Tissue homogenates were serially diluted and plated on PIA and CFU determination was made 18 hours later. Final results were expressed as CFU/ml for nasal washes and CFU/g for other organ tissues.

### Opsonophagocytic killing assay

Luminescent opsonophagocytosis assay was performed according to the method as described (20). Immune sera from mice vaccinated with 10^7^ or 10^9^ CFU of SL3261 (pLAFRO9) were used, and 3-fold serial dilutions ranging from 1:100 – 1:59,049 were tested in triplicates. Sera collected from vector- and PBS treated mice were also tested using the same dilutions. Constitutively bioluminescent *P. aeruginosa* PAO9 strain was constructed by immobilizing the plasmid pUC18miniTn7T-*lux*-Tp into the wild-type PAO9 strain to generate PAO9 P1*-lux* according to the method described (21). Briefly, 25 μl of each opsonophagocytosis component: *P. aeruginosa* PAO9 P1-*lux*, from log phase cultures diluted to 2 x 10^6^ CFU/ml; diluted baby rabbit serum (1:10); 2 x 10^7^ polymorphonuclear leukocyte (PMN); and sera collected from PBS-, vector-, or vaccine-immunized mice. The percentage of killing was determined by comparing the relative luciferase units (RLU) derived from assays lacking serum to the RLU obtained from assays with vector or vaccine sera. The assay was performed in 96-well plates, following a 120 minute-incubation at 37°C shaking at 250 RPM. Microtiter plates were read using an Envision Multilabel plate reader (PerkinElmer).

### Statistical analysis

All analyses were performed using GraphPad Prism version 6 software. ELISA endpoint titers were calculated using the linear regression of duplicate measurements of adjusted OD_405_ and were expressed as the reciprocal dilution. The x-intercept served as the endpoint titer. Antibody titers were compared using the Kruskal-Wallis test for comparison of three groups or the Mann-Whitney U test for two group analysis. The results of survival studies were represented using Kaplan-Meier survival curves and were analyzed by the log-rank test.

## RESULTS

### *S. typhimurium* SL3261 containing pLAFRO9 expresses *P. aeruginosa* serogroup O9 LPS

We cloned the O antigen locus from the *P. aeruginosa* serogroup O9 strain, PAO9, into the broad host range cosmid, pLAFR376, using standard techniques. The plasmid was transferred to *S. typhimurium* SL3261 using P22 transduction. To confirm expression of *P. aeruginosa* serogroup O9, LPS was purified from PAO9, SL3261, and SL3261 (pLAFRO9), and separated by SDS-PAGE. A silver stain gel of the LPS fragments is shown in **Fig. 1A**. The pattern of banding is more similar between the two SL3261 samples (lane 2 and lane 3) and distinct from PAO9 LPS (lane 1), however banding is seen in all lanes, indicative of LPS isolation. The LPS was analyzed by immunoblotting and reactivity was detected with polyclonal anti-sera specific for serogroup O9 (**Fig. 1B**). Sera reacted with LPS from PAO9 (lane 1) and with LPS from the *Salmonella* containing pLAFRO9 (lane 3), no reactivity to LPS from SL3261 (lane 2) was seen. These data confirm expression of the *P. aeruginosa* serogroup O9 LPS in our vaccine strain.

**Figure 1.**
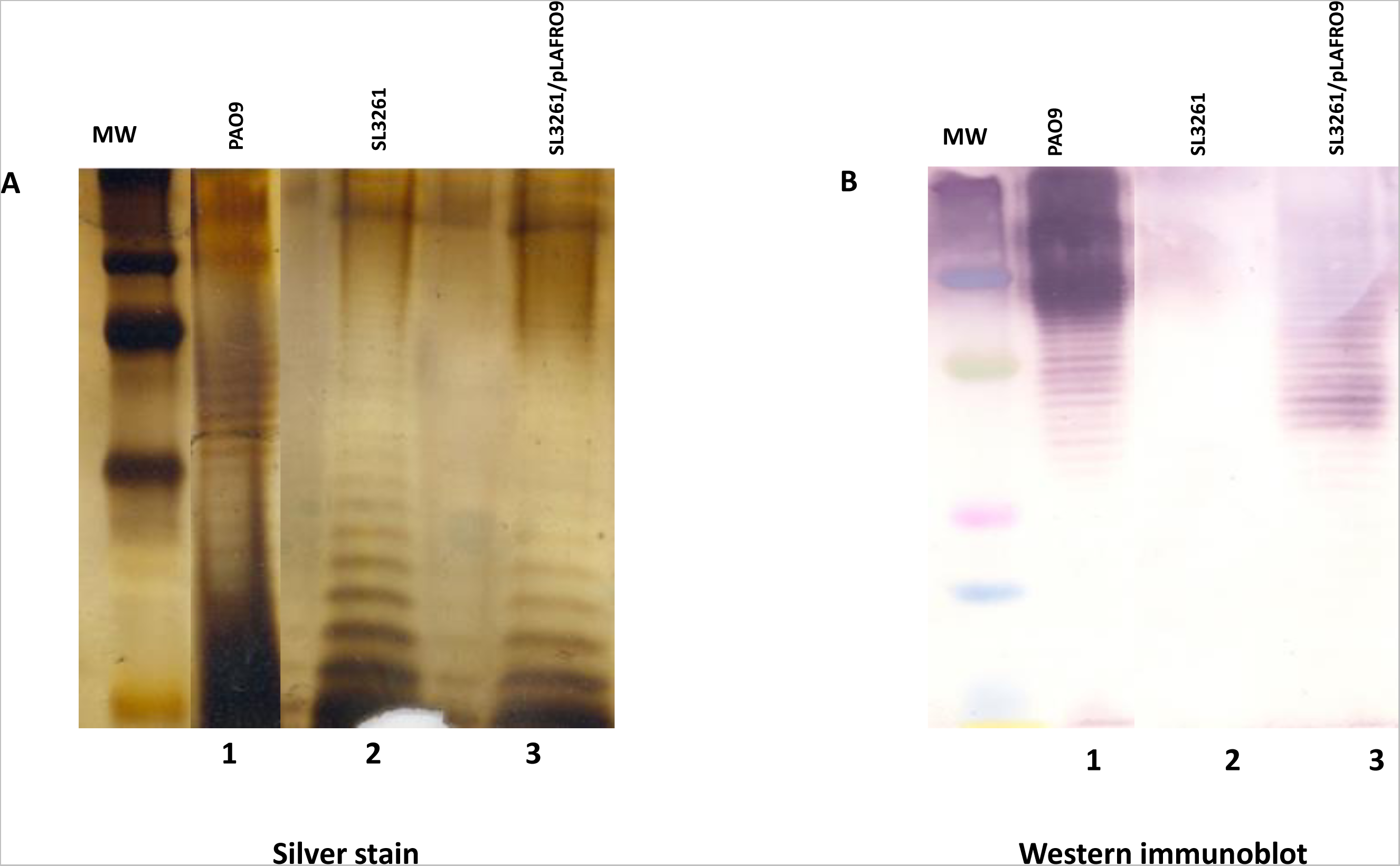
Expression of *P. aeruginosa* serogroup O9 in *S. typhimurium*. LPS was extracted, applied to SDS-PAGE, and visualized with silver stain (A) or by (B) immunoblot analysis using *Pseudomonas* serotype O9-specific rabbit polyclonal antibody, followed by incubation with anti-rabbit secondary antibody conjugated to alkaline-phosphatase (Sigma). *P. aeruginosa* serogroup O9 strain PAO9 (Lane 1), *S. typhimurium* SL321 (lane 2), and from *S. typhimurium* SL3261 containing the plasmid expressing the *P. aeruginosa* serogroup O9 antigen (pLAFRO9) (Lane 3). Molecular weight (MW).

### Intranasal immunization of mice with SL3261(pLAFRO9) induces serum antibody responses that confers protection in an acute pneumonia model

Our group has previously demonstrated the safety and efficacy of SL3261 expressing PA103 serogroup O11 O-antigen (18, 22). Our previous studies with *P. aeruginosa* serogroup O11 expression in SL3261 showed intranasal administration of 10^7^ CFU/mouse elicited a protective immune response (18). Based on these prior observations, we intranasally immunized mice on day 0 and day 14 with 10^7^ CFU/mouse of the vaccine (SL3261 (pLAFRO9)) or the vector control (SL3261 (pLAFR376)). The kinetics of the *Pseudomonas*-specific serum IgG and IgM antibody responses in sera collected from vaccine-immunized mice were compared to those immunized with PBS- or vector-only. We compared *Pseudomonas*-specific antibody response on day 14 and on day 28, two weeks after the booster dose.

Interestingly, intranasal immunization using 10^7^ CFU/mouse failed to generate a serogroup O9-specific antibody response. We did not observe a substantial difference in serum IgM response in the immunized mice compared to the vector- or PBS-treated mice (**Fig. 2A**). A similar trend was also observed with the serogroup O9 IgG response (**Fig. 2B**). Notably, we did not observe a substantial increase in the immune response in any of immunized mice on day 28, after receiving the booster dose.

**Figure 2.**
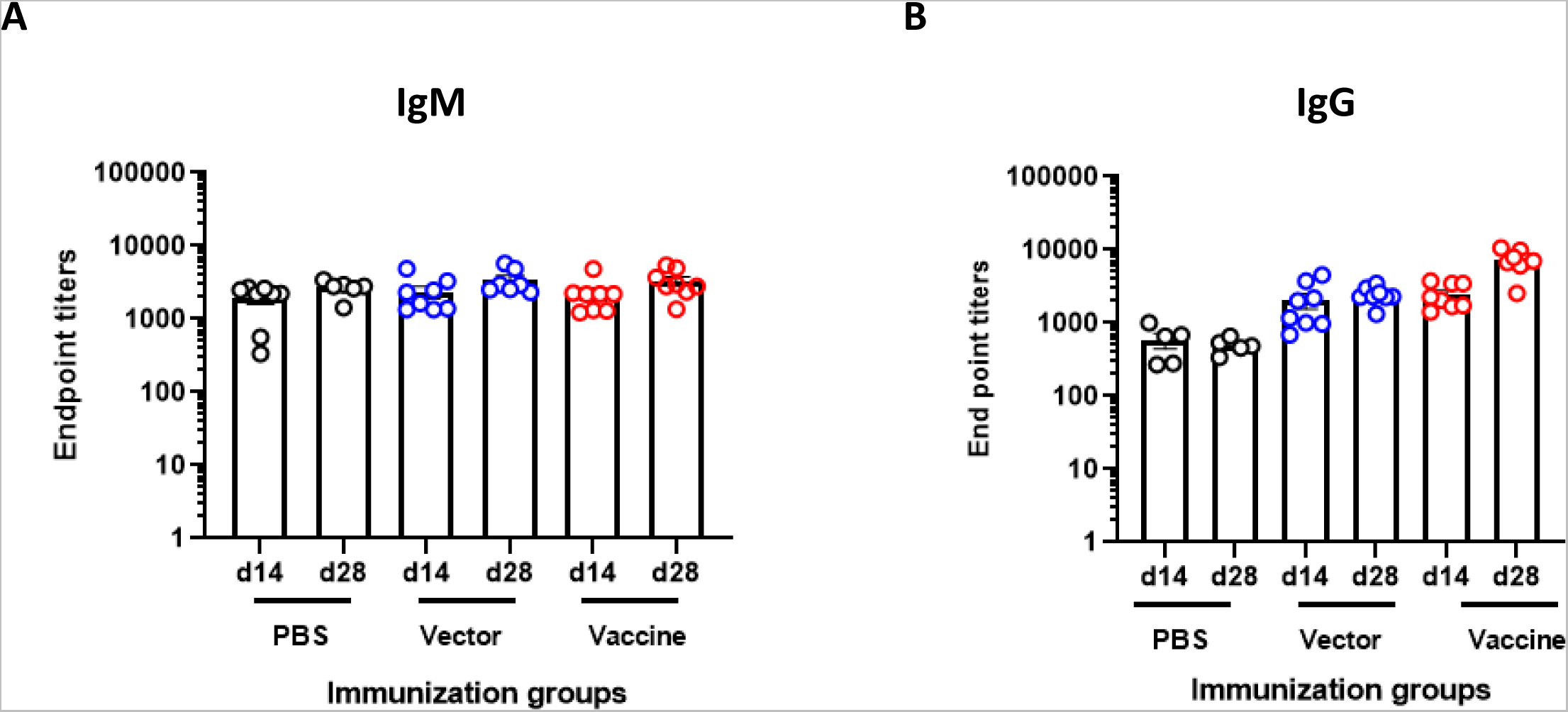
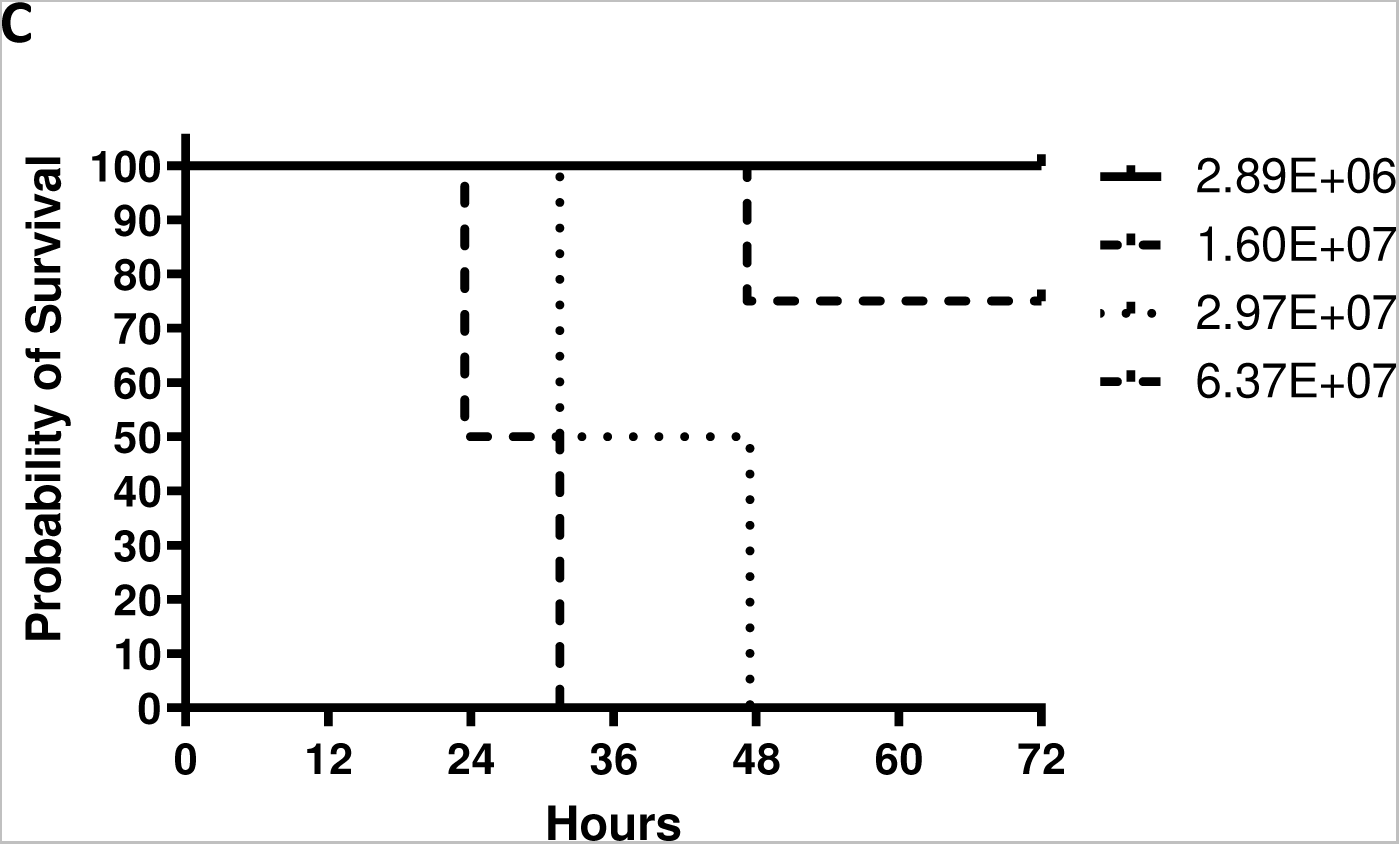
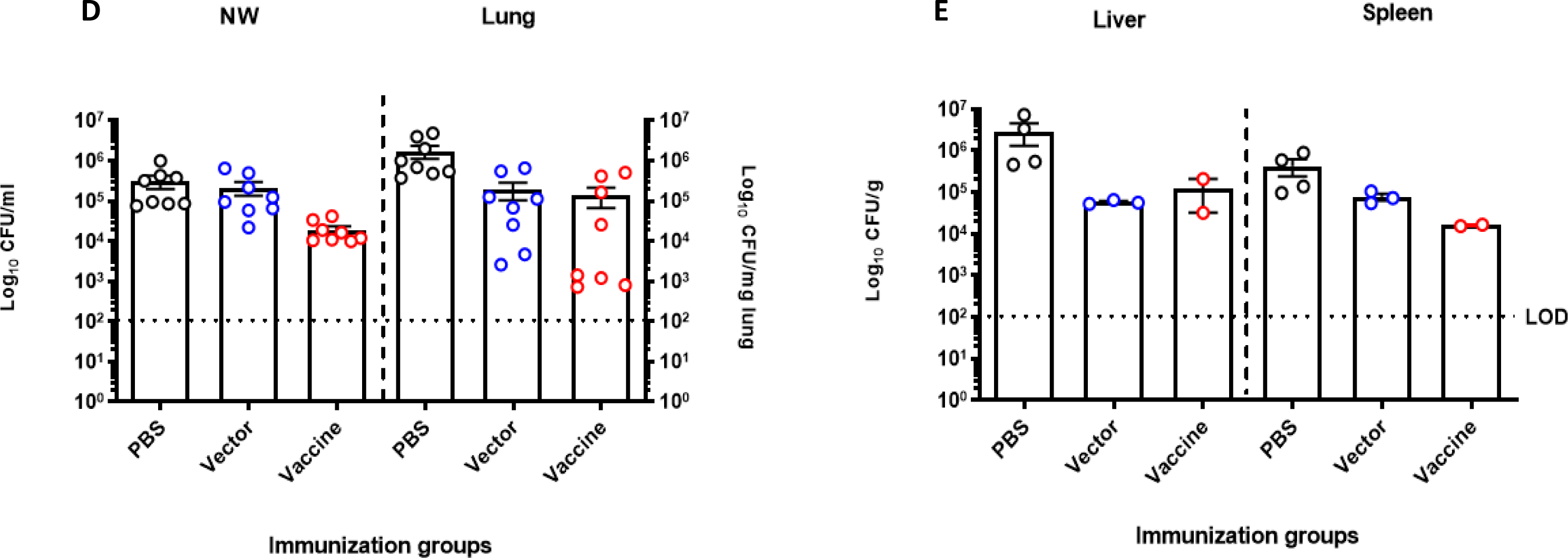
**Serum antibody response of BALB/c mice following intranasal immunization with *S. typhimurium* SL3261 expressing *P. aeruginosa* serogroup O9 and bacterial loads in organs of intranasally immunized BALB/c mice after intranasal lethal challenge with *P. aeruginosa* PAO9**. Mice were immunized intranasally on day 0 and day 14 with 10^7^ CFU/mouse of the vector control (SL3261 (pLAFR376)) or the vaccine (SL3261 (pLAFRO9)). Sera were collected 2- and 4-weeks post-vaccination, and the data were analyzed by the Mann-Whitney *U* test. (A) Serum IgM and (B) IgG response to *P. aeruginosa* PAO9 whole antigen. (C) Survival rates for naïve mice after intranasal challenge with various doses of *P. aeruginosa* PAO9 (n=4 mice/group). (D) Bacterial load in the nasal wash and lungs, (E) liver and spleens of intranasally immunized mice 24 hours post-challenge with 2 x 10^7^ CFU of PAO9. All samples were plated for viable CFU on Pseudomonas isolation agar (PIA). Each point represents a single mouse.

To examine the ability of the vaccine strain to confer protection against acute pneumonia in intranasally immunized animals, mice were challenged by intranasal infection with *P. aeruginosa* PAO9. Prior to this, we tested the virulence of PAO9 delivered via the intranasal route in 20 µl. As shown in **Fig. 2C**, increased doses decreased the time to when animals became moribund or were under distress, with a 50% lethal dose (LD_50_) calculated for this strain to be ∼2.0 x 10^7^ CFU, according to Reed and Muench (23). Immunized mice were challenged with ∼1X the LD_50_ (2.0 x 10^7^ CFU) and mice were euthanized at 24 hours post-infection to assess colonization in the upper (nasal wash) and lower (lung) respiratory tract. Livers and spleens were also collected for the determination of bacterial CFU. As seen in **Fig. 2D**, no statistical difference was seen in bacterial loads in the lungs of vaccine-immunized animals and vector-immunized animals, but both were different compared to PBS-immunized mice. Furthermore, dissemination to liver and spleens of immunized mice was also detected. There was a significant decrease in the bacterial load in the liver and spleen in the vaccine immunized mice relative to the CFUs recovered from PBS immunized mice. However, no statistical difference in bacterial CFU was observed between mice immunized with the vaccine (SL3261 (pLAFRO9) and mice inoculated with the vector (SL3261 (pLAFR376)) (**Fig. 2E**).

As we analyzed the ELISA data from this initial experiment, we noted that the IgG levels were not as robust as what we had previously observed for serogroup O11 (18), suggesting that higher doses of the vaccine might be needed.

We subsequently modified our vaccination protocol and immunized mice with different amounts of the vector or vaccine; we administered 10^7^ CFU on day 0 and 14, 10^7^ CFU at day 0 and 10^9^ CFU at day 14, or 10^9^ CFU at day 0 and 10^9^ CFU at day 14. Serum samples were taken at 4 weeks post-vaccination, and we observed a direct correlation between the immunization dose and the level of O9-specific IgG, but not IgM (**Fig. 3A & 3B**), with the mice administered 10^9^ CFU of SL3261 (pLAFRO9) exhibiting significantly increased serogroup O9-specific IgG response after the booster immunization (**Fig. 3B**).

**Figure 3.**
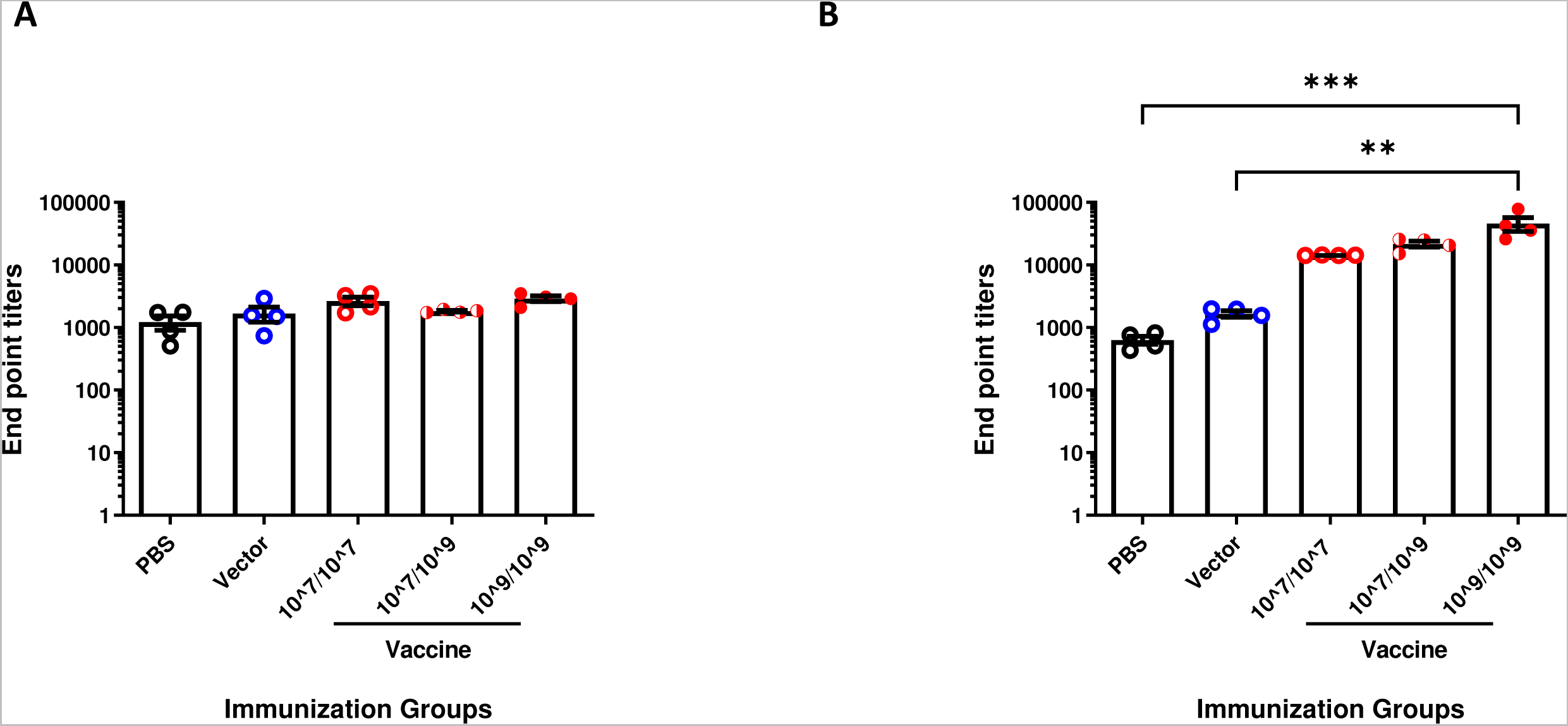
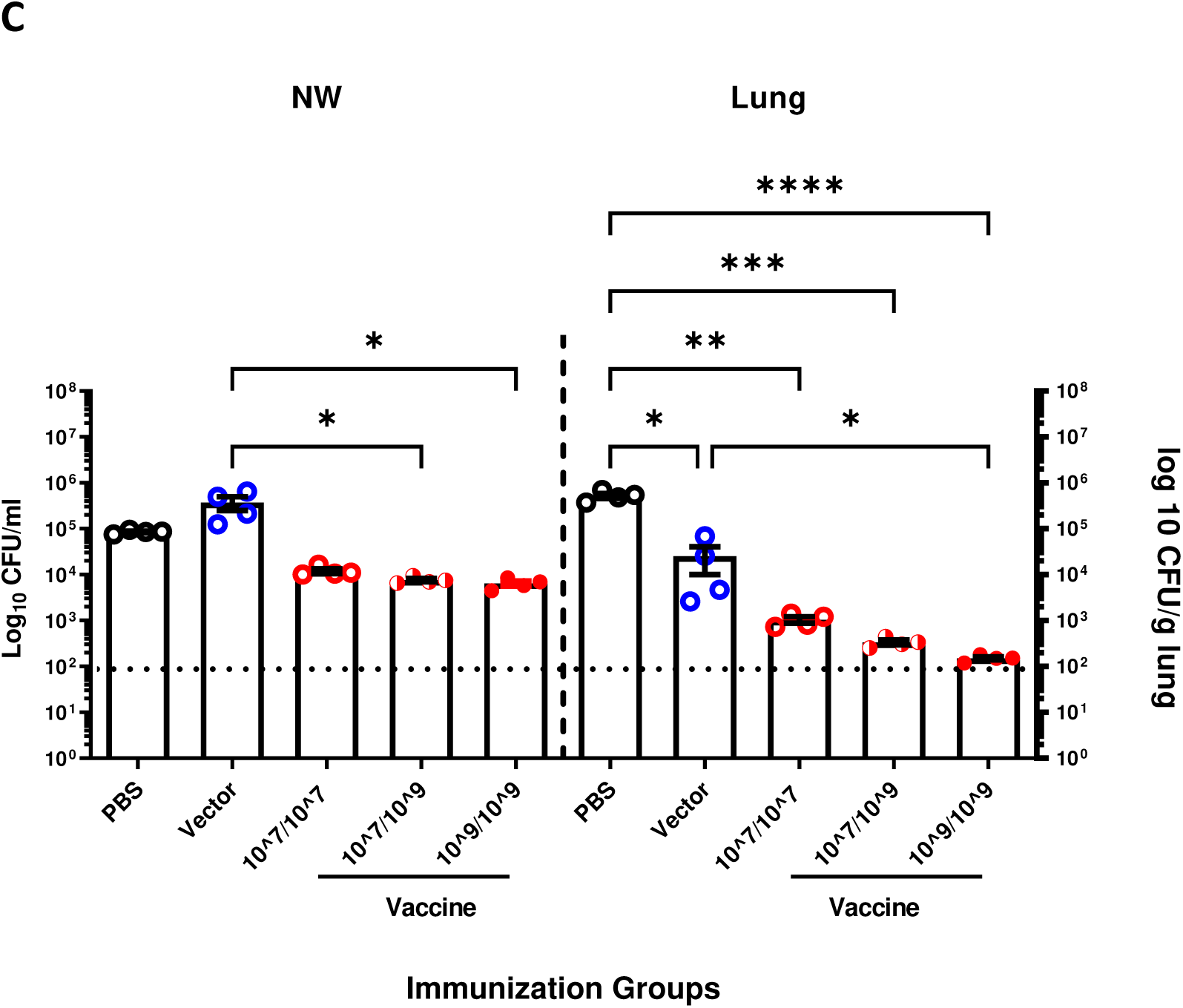
Immune response and bacterial burden in response to immunizing BALB/c mice with various doses of *S. typhimurium* SL3261 expressing *P. aeruginosa* serogroup O9. Anti-*P. aeruginosa* serogroup O9 serum (A) IgM and (B) IgG antibody response after intranasal vaccination of BALB/c mice to various doses of vector or vaccine. (C) Bacterial load in the nasal wash and lungs of intranasally immunized mice 24 hours post-challenge with 4.5 x 10^7^ CFU of PAO9. All samples were plated for viable CFU on PIA. Each point represents a single mouse. Data were analyzed by one-way ANOVA, * P<0.05, **P<0.01, ***P<0.001, ****P<0.0001. Error bars represent the mean and SEM.

At 6 weeks post-vaccination, mice were challenged with 4.5 x 10^7^ CFU of PAO9 (∼2.5X the LD_50_). After 24 hours, the bacterial colonization in the upper and lower respiratory tract was determined. There was an inverse correlation between the CFU of PAO9 in either upper or lower respiratory tract and the level of IgG (**Fig. 3C**). Notably, we observed a strong correlation between the level of IgG and the subsequent clearance from nasal washes and the lungs, as the bacterial burden tended to decrease with increasing doses of SL3261 (pLAFRO9) administered.

Based on the results from the previous experiment, we repeated the immunization by administering 10^9^ CFU of SL3261 (pLAFRO9) vaccine at day 0 and boosted each mouse with the same dose, 10^9^ CFU on day 14. ELISA analysis using sera collected four weeks post-immunization from BALB/c mice that received the vaccine revealed robust *P. aeruginosa* serogroup O9 LPS-specific IgM and IgG antibody response when compared with sera from the vector-immunized or PBS control mice (**Fig. 4A & 4B**).

**Figure 4.**
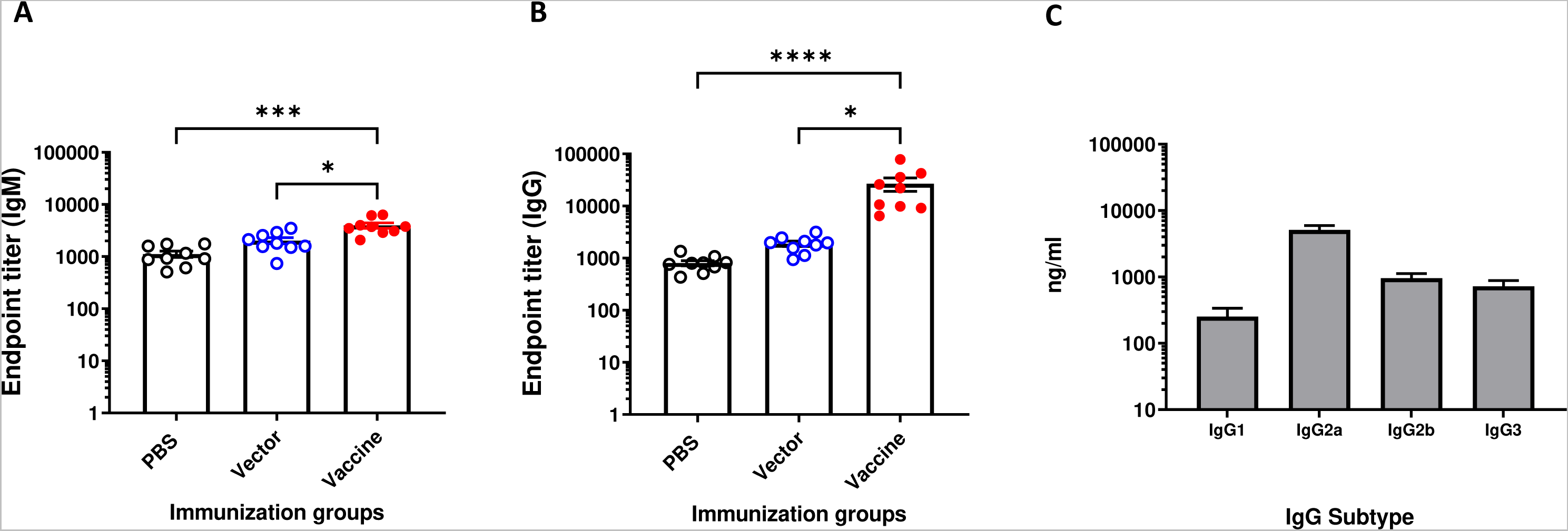
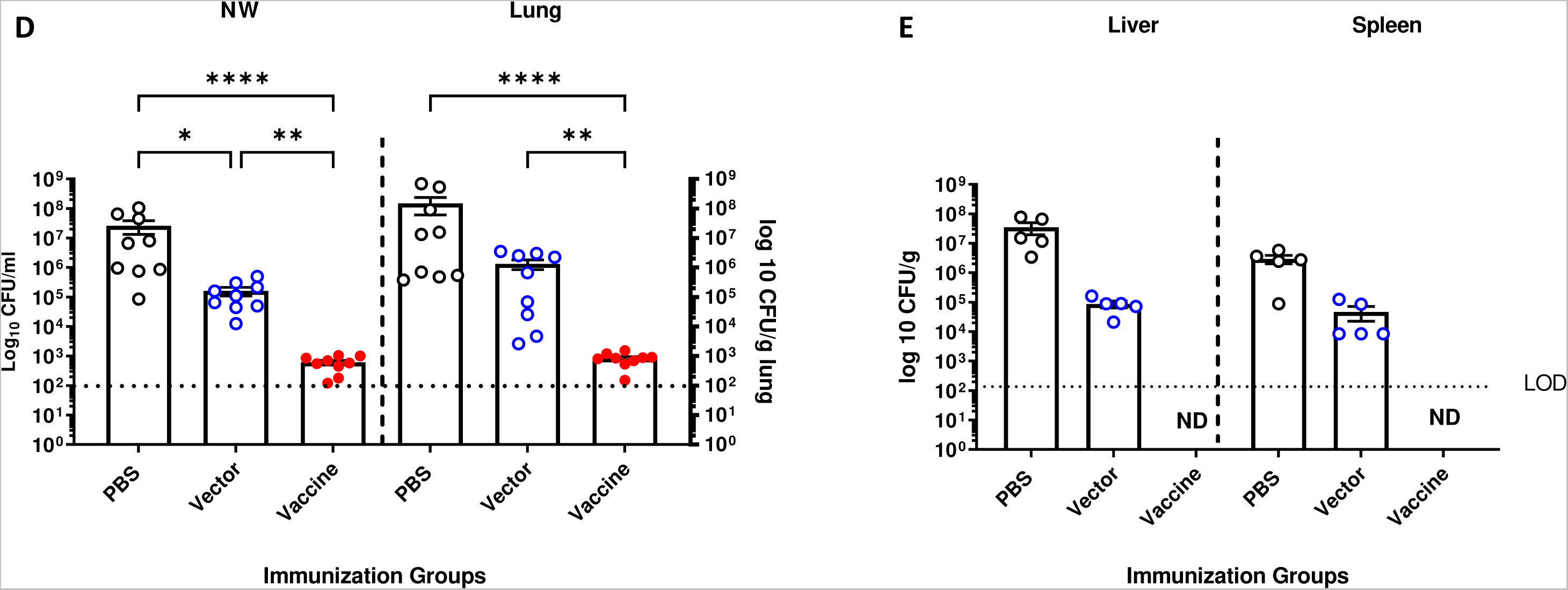
Immune response and bacterial burden in response to immunizing BALB/c mice with *S. typhimurium* SL3261 expressing *P. aeruginosa* serogroup O9 at 10^9^ CFU/mouse. Anti-*P. aeruginosa* serogroup O9 serum (A) IgM, (B) IgG antibody, and (C) IgG isotype response after intranasal vaccination of BALB/c mice with 10^9^ CFU/mouse of SL3261 (pLAFR376) (vector) and SL3261 (pLAFRO9) (vaccine). (D) Bacterial load in the nasal wash and lungs, and (E) liver and spleens of intranasally immunized mice 24 hours post-challenge with 8.6 x 10^7^ CFU of PAO9. All samples were plated for viable CFU on PIA. Each point represents a single mouse. Data were analyzed by one-way ANOVA. * P<0.05, **P<0.01, ***P<0.001, ****P<0.0001. Error bars represent the mean and SEM.

The IgG subtype responses to *P. aeruginosa* PAO9 were determined for vaccine-immunized mice. Intranasal immunization elicited significantly higher levels of IgG2a, IgG2b and IgG3. The levels of IgG2a antibodies were significantly higher compared to IgG1 and IgG3 (*P<0.001*) (**Fig. 4C**).

Given that we observed a significant increase in *P. aeruginosa* Ig titers following intranasal immunization, we next sought to determine the level of protection against PAO9 challenge in immunized mice. At 6 weeks post-vaccination, mice were challenged with 8.6 x 10^7^ CFU of PAO9 (∼4X the LD_50_). Again, we observed a statistically significant difference in the bacterial counts recovered from vaccine-immunized mice when compared to the PBS and vector-immunized controls in either upper (nasal wash) or lower (lung) respiratory tract (**Fig. 4D**). Furthermore, no bacteria were recovered from the spleen nor liver tissues from vaccine-immunized mice (**Fig. 4E**).

### Antibody response induced by intranasal vaccination mediates opsonic killing of *P. aeruginosa* PAO9 *in vitro*

We previously demonstrated immunization of *Salmonella* carrying pLPS2 results in the production of O11 O-antigen-specific antibodies and induces opsonic antibodies (18). Here, we examined the efficiency of sera collected from mice immunized and boosted with 10^7^ or 10^9^ CFU of the vaccine to promote opsonization and phagocytosis of *P. aeruginosa* PAO9 by PMNs. As a control, we performed the assay in absence of vaccine-immune sera to further confirm the role of antibodies in protection. Pooled antisera from vaccine-immunized mice mediated high-level opsonophagocytic killing (>50%) of PAO9 strain at serum dilutions up to 1:2,700 compared to the limited killing observed with pooled sera from vector- and PBS-immunized mice (**Fig. 5**).

**Figure 5.**
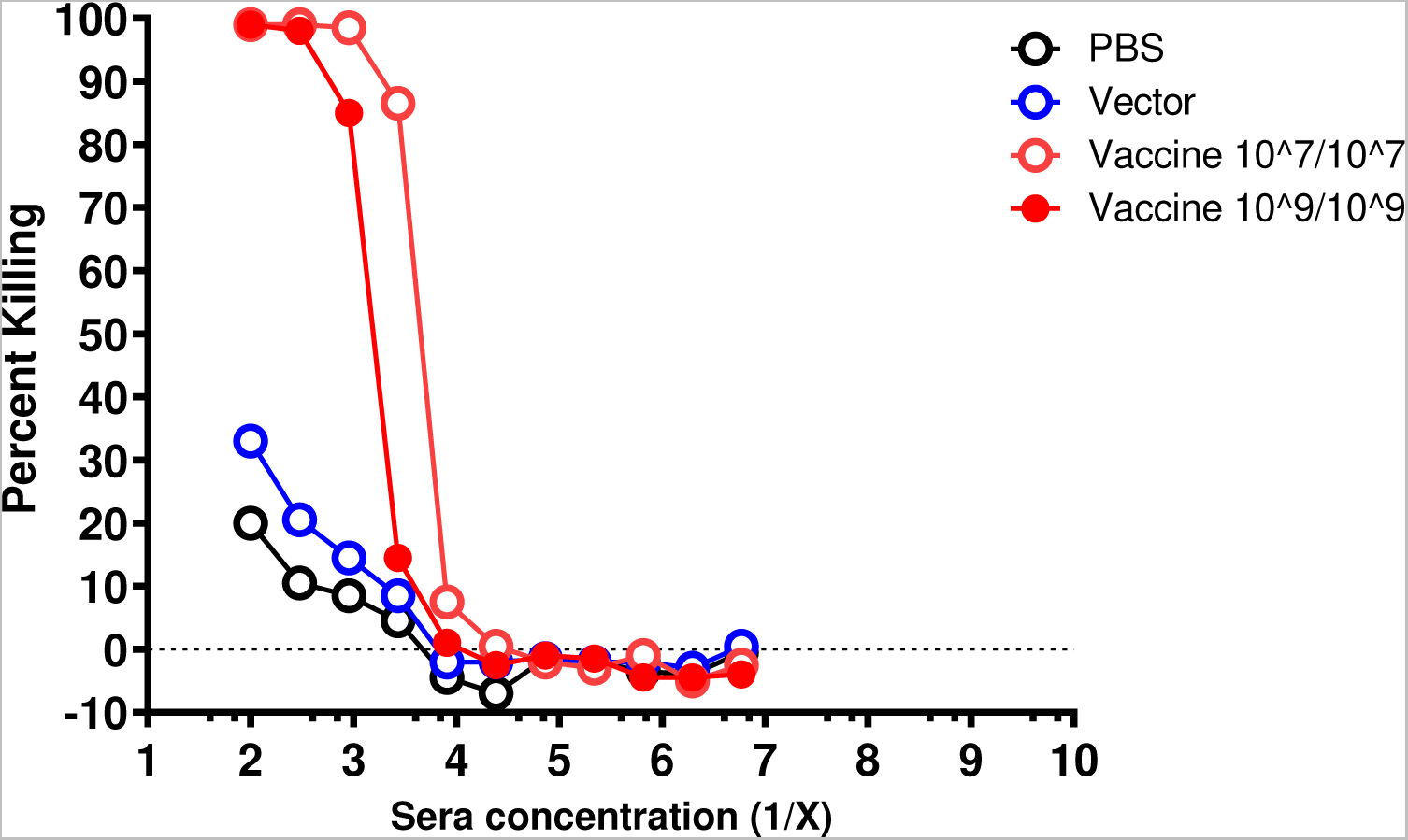
Antibody response induced by intranasal vaccination mediates efficient opsonic killing of *P. aeruginosa* PAO9 *in vitro.* Opsonophagocytic killing of *P. aeruginosa* PAO9 P1-*lux* using dilutions of pooled antisera collected from intranasally PBS-, SL3261 (pLAFR376) (vector)-, and SL3261 (pLAFRO9) (vaccine)-immunized BALB/c mice. Plates were read at 120 minutes following the co-incubation of the opsonophagocytosis assay components.

### Passive transfer of antisera from immunized animals to naïve mice provides protection from acute *P. aeruginosa* pneumonia

To identify whether mice could be protected from pneumonia caused by infection with *P. aeruginosa*, undiluted antisera from PBS-, vector-, or vaccine-immunized mice were transferred intranasally to naive BALB/c mice at the same time that they were infected with a sublethal dose of PAO9. Briefly, female BALB/c mice 8-10 weeks old were divided into 3 groups, 5 mice each. Each group received 10 µl (5µl/nostril) of sera collected from mice immunized with: PBS, 10^7^ CFU (SL3261 (pLAFR376)), or 10^9^ CFU (SL3261 (pLAFRO9)). Immediately after serum administration, mice were intranasally infected with 1.1 x 10^7^ CFU/mouse (∼0.5X the LD_50_) of *P. aeruginosa* PAO9. Mice were euthanized 24 hours post-infection, and the nasal wash, lung, liver, and spleen from each mouse was removed and homogenized to assess bacterial counts.

The number of viable bacteria recovered from the nasal wash and lung tissue of mice receiving vaccine-immune sera was shown to drop significantly as compared to those recovered from mice receiving PBS- and vector-immune sera. We observed a statistically significant difference in the bacterial counts recovered from mice receiving antisera from vaccine-immunized mice when compared to those treated with antisera from PBS- and vector-immunized controls (**Fig. 6A & 6B**) indicating that the immune sera provide protection from infection.

**Figure 6.**
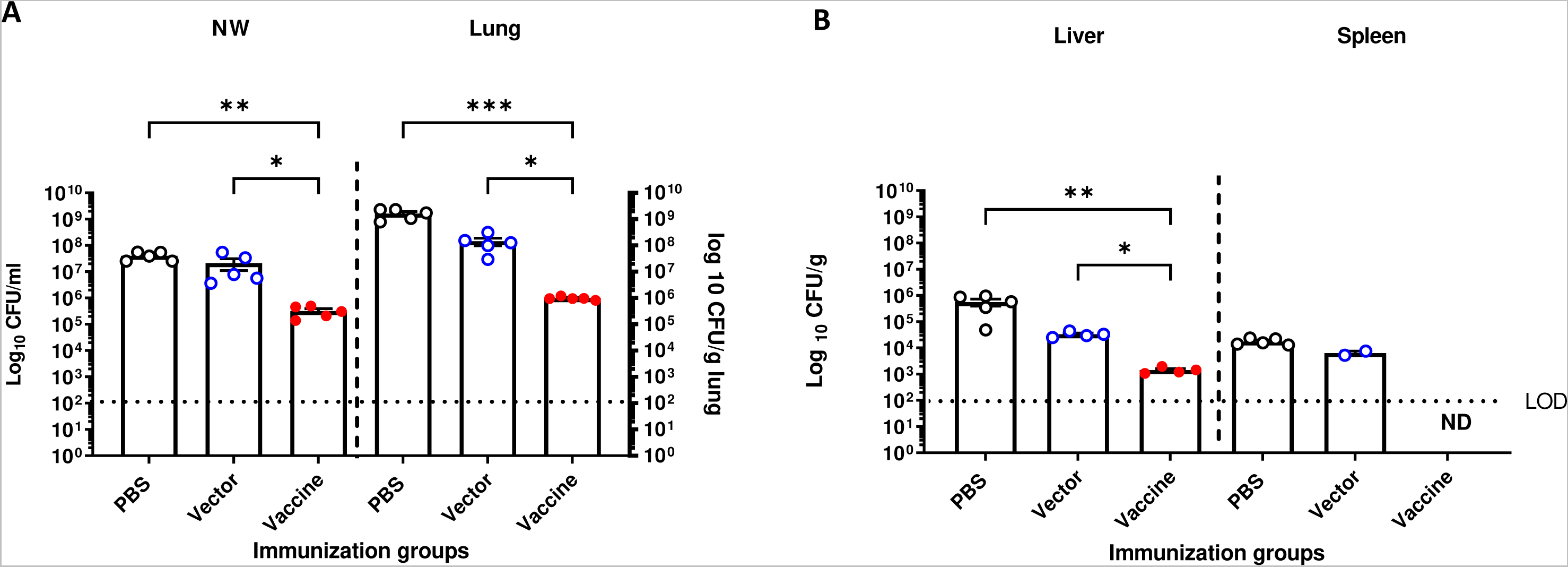
Passive antisera transfer from immunized animals provides protection to acute *P. aeruginosa* pneumonia in naïve mice. Bacterial loads in (A) nasal wash and lungs, (B) liver and spleens of BALB/c mice after passive intranasally transferred antisera immediately followed by challenge with 1 x 10^7^ CFU of PAO9. Mice were euthanized 24 hours post-infection. All samples were plated for viable CFU on PIA. Each point represents a single mouse. Data were analyzed by one-way ANOVA. * P<0.05, **P<0.01, ****P<0.001. Each point represents a single mouse. Error bars represent the mean and SEM.

## DISCUSSION

Lipopolysaccharide (LPS) is an immunodominant protective antigen for *P. aeruginosa*. The O antigen portion of *P. aeruginosa* LPS confers serogroup specificity and *P. aeruginosa* has 20 different serogroups that vary by sugar composition (24), however 10 are most commonly associated with infection (9–12).

Several groups have attempted to develop vaccines to prevent infection based on LPS and these have met with limited success; there is currently no *P. aeruginosa* licensed vaccine (25, 26). Recently, Nasrin et al. (12) reported a “Multiple Antigen Presenting System” for *P. aeruginosa* based on high molecular weight polysaccharides (17) and have targeted 8 of the most common O antigen serogroups (12), however serogroup O8 and O9 are missing from this system. Both serogroup O8 and O9 are acid labile in nature and as such have never been included in a conjugate vaccine cocktail (27).

Our laboratory previously characterized a vaccine that confers serotype-specific protection against *P. aeruginosa* challenge (18). The vaccine consists of *Salmonella enterica* serovar Typhimurium strain SL3261, an attenuated *aroA* mutant (28), expressing the entire O-antigen locus from a *P. aeruginosa* serogroup O11 strain (19). The *aroA* gene encodes 3-enolpyruvyl-shikimate-5-phosphate synthetase, an enzyme required for the synthesis of amino acids and growth. Intranasal vaccination with this *Salmonella* strain conferred complete protection in mice with challenge doses of 5X the LD_50_ of both cytotoxic and noncytotoxic *P. aeruginosa* serogroup O11 strains. Moreover, administration of antibodies from vaccinated mice directly into the nasal passageway and lungs of infected mice was able to confer protection when administered up to six hours after infection.

A caveat to this vaccine is that protection is only directed to serogroup O11 strains. *P. aeruginosa* serogroup O9 is another serogroup commonly found in infection, although it has been “neglected” in these previous *P. aeruginosa* vaccine cocktails. Studies of Faure et al. (9) noted in a survey of clinical isolates that 4% (4/99 total) were serogroup O9 (2% in chronic infections and 2% in acute infections), but that these were not associated with mortality. Using *in silico* serotyping, Thrane et al. (10) found 1.25% of the 1120 genomes they surveyed were serogroup O9. Interestingly, when looking specifically at genomes of isolates from cystic fibrosis patients, this number was much lower (0.38% from 529 genomes) (10). Ozer et al. (11) found serogroup O9 strains accounted for 2% of the isolates they sequenced from diverse sources. And more recently Nasrin et al. (12) surveyed 413 invasive *P. aeruginosa* from 10 different countries worldwide and found ∼3% of them were serogroup O9.

Using a similar approach to what we previously reported expression of the *P. aeruginosa* serogroup O11 O antigen locus in *Salmonella enterica* serovar Typhimurium (19), here we cloned the serogroup O9 locus. We showed that the recombinant *Salmonella* strain expressed *P. aeruginosa* serogroup O9 LPS (**Fig. 1**). We extracted DNA and performed DNA sequence analysis and confirmed the sequence corresponded to the serogroup O9 locus as reported by Raymond et al. (29).

We noted a number of subtle but interesting differences between our original experiments with serogroup O11 (18) and those performed here with serogroup O9. We needed to modify our vaccination protocol and increase the dose of *Salmonella* to 10^9^ CFU to elicit a robust and protective immune response. This may have reflected low expression of the plasmid-borne genes of the serogroup O9 locus, reduced amount of LPS expressed on *Salmonella*, and/or the lability of the O antigen itself. However, once the inoculum was optimized to generate an immune response, it was protective.

Prior to the initiation of immunization studies, we needed to find a strain to assess protection. We had not been able to find any reports of using serogroup O9 strains in models of infection, therefore we determined the virulence of PAO9 in our murine intranasal acute pneumonia model. Interestingly, we noted that this strain was not very pathogenic and therefore we needed to give large doses to mice to monitor vaccine-mediated protection. This was also the case for additional serogroup O9 strains obtained from collaborators. Supporting this, Ozer et al. noted that the majority of the serogroup O9 isolates that they characterized (14/15) were in Group A and most Group A strains were lacking the *exoU* gene (11). Similarly, Faure et al. determined that only 1 of the 4 serogroup O9 clinical isolates that they examined secreted ExoU (9). ExoU is a marker for highly virulent strains, especially associated with acute lung infections (30), thus if serogroup O9 strains we tested were lacking *exoU* it could explain why they are less able to cause severe infections.

To our knowledge, there has only been one report of a serogroup O9-specific epidemic, which was an outbreak of dermatitis in a whirlpool at a hotel in Atlanta, Georgia in 1981 (31). Whether the lack of detection of outbreaks caused by serogroup O9 isolates is because of avirulence due to the lack of *exoU* and/or the acid labile nature of the O antigen itself is not known but is tempting to speculate. However, that is not to assume that all serogroup O9 *P. aeruginosa* isolates are innocuous. In 2022, there was a fatal case of community-acquired *P. aeruginosa* pneumonia in an otherwise immunocompetent individual following SARS-CoV-2 infection (32).

While many of the current available bacterial vaccines are based on either purified polysaccharides (pneumococcal polysaccharide vaccine) or polysaccharide-protein conjugates (pneumococcal conjugate vaccines, meningococcal conjugate [menACWY] vaccines, and Hib vaccines). Using recombinant attenuated *Salmonella* for heterologous expression has the advantage of ease of expression of polysaccharide antigens and generating an immune response more typical of a polysaccharide-protein conjugate (33). Supporting this, we found that our vaccine induced high levels of IgG2a, IgG2b, and IgG3 to serogroup O9 (**Fig. 4C**).

We have previously shown that we could express *P. aeruginosa* serogroup O11 on the human licensed typhoid vaccine strain, *Salmonella* Typhi Ty21a (34), when it contained the plasmid pLPS2 (35) suggesting the feasibility of this approach for human use. Since that time, methods for stable integration of O antigen genes have been developed (36) as well as techniques for the expression of multiple serogroups in the same *S.* Typhi Ty21a strain (37).

We have also found that vaccination with SL3261 (pLAFRO9) induced antibodies that were capable of mediating opsonophagocytosis (**Fig. 5**) and we could transfer protection from immunized mice to naïve mice with the sera alone (**Fig. 6**). This suggests the possibility that this vaccination protocol could be used to develop a passive immunotherapy specific for the “neglected” *P. aeruginosa* serogroup O9. Such an approach has been used for *P. aeruginosa* serogroup O11. There is a fully human anti-LPS IgM monoclonal antibody (Panobacumab) that has been tested in patients with nosocomial *Pseudomonas* pneumonia due to serogroup O11. Treatment with Panobacumab resulted a shorter time to clinical resolution compared to untreated patients (38), but this is only effective for patients infected with a serogroup O11 *P. aeruginosa* strain. The advantage is that passive immunotherapy would be applicable to all patients, including the immunocompromised who cannot mount an immune response and who are particularly at risk for *P. aeruginosa* hospital infections. The long-term goal of this research is ultimately the development of a cocktail including all prominent serogroups that could be given to people to protect them against *P. aeruginosa* infection by any of the typically encountered strains.

## Acknowledgements

This work was supported by grants to J.B.G. from the National Institutes of Health (1 R01 AI50230, 1 R01 AI68112, and 1 R21 AI53842) and from the Cystic Fibrosis Foundation (GOLDBE00G0 and GOLDBE05P0). J.M.S. and M.R.D were supported in part by the National Institutes of Health through the University of Virginia Infectious Diseases Training Grant AI07406.

